# An intrinsically Disordered RNA Binding Protein Modulates mRNA Translation and Storage

**DOI:** 10.1101/2023.05.22.541713

**Authors:** Mashiat N. Chowdhury, Xin Chen, Hong Jin

## Abstract

Many proteins with intrinsically disordered regions interact with cytoplasmic ribosomes. However, many of the molecular functions related to these interactions are unclear. In this study, using an abundant RNA-binding protein with a structurally well-defined RNA recognition motif and an intrinsically disordered RGG domain as a model system, we investigated how this protein modulates mRNA storage and translation. Using genomic and molecular approaches, we show that the presence of Sbp1 slows ribosome movement on cellular mRNAs and promotes polysome stalling. Sbp1-associated polysomes display a ring-shaped structure in addition to a beads-on-string morphology visualized under electron microscope. Moreover, post-translational modifications at the RGG motif play important roles in directing cellular mRNAs to either translation or storage. Finally, binding of Sbp1 to the 5’UTRs of mRNAs represses both cap-dependent and cap-independent translation initiation of proteins functionally important for general protein synthesis in the cell. Taken together, our study demonstrates an intrinsically disordered RNA binding protein regulates mRNA translation and storage via distinctive mechanisms under physiological conditions and establishes a framework with which functions of important RGG-proteins can be investigated and defined.

## Introduction

RNA-binding proteins play critical and diverse roles in regulating post-transcriptional gene expression (Gebauer et al., 2021; Gerstberger et al., 2014; Hentze et al., 2018). This family of proteins often contains one or more RNA binding domains (RBDs) that sequence-specifically or structure-specifically bind RNAs. Many RNA-binding proteins are localized in both the nucleus and the cytoplasm, where they intimately regulate a wide range of nuclear and cytoplasmic processes pertinent to RNA metabolism including RNA-specific epigenetic control, RNA splicing, transport, translation, stability and degradation.

As an integral step of post-transcriptional gene regulation, translational regulation allows the cell to rapidly, often reversibly, respond to environmental perturbations and internal cues, fine-tuning protein synthesis for proper cellular homeostasis and physiology (Jackson et al., 2010; Sonenberg and Hinnebusch, 2009). Translation is regulated at each step of protein synthesis including initiation, elongation, termination and recycling, wherein RNA binding proteins play critical roles in controlling global and the transcript-specific translation.

RGG (Arginine-Glycine-Glycine)-motif containing proteins are the second most common of the known RBPs in the human genome (Ozdilek et al., 2017; Thandapani et al., 2013). This family of proteins is broadly defined as proteins with closely spaced multiple RGG or RG repeats, interspersed with various amino acids (Chowdhury and Jin, 2022; Thandapani et al., 2013). In addition to varying numbers of the RGG repeats, the length of spacing and the identity of amino acids between the RGG repeats differ as well, contributing to the compositional complexity of RGG-proteins.

The unique combination of the arginine and glycine residues enables RGG-proteins to form specific, yet adaptable and versatile molecular interactions with a broad range of proteins and nucleic acids. Furthermore, post-translational modifications at the arginine within the RGG motif extends the capacity of RGG-proteins for fine-tuned interaction and regulation. As such, RGG proteins are functionally involved in a wide range of essential cellular processes from DNA repair (Déry et al., 2008; Mastrocola et al., 2013; Yu et al., 2012), chromatin structure modulation (Erard et al., 1988; Yan et al., 2021), transcription (Mowen et al., 2004; Rickards et al., 2007), pre-mRNA splicing (Lee et al., 2018; Zhou et al., 2019), RNA transport (Singleton et al., 1995), translation (Athar and Joseph, 2020; Chen et al., 2014) to ribonucleoprotein biogenesis (Ginisty et al., 1998).

Moreover, the intrinsically disordered nature of the RGG motif enables the protein to phase-separate into membraneless granules such as cytoplasmic processing bodies (P bodies) and stress granules, shuttling the targeted mRNAs away from the translation pool (Bourgeois et al., 2020; Lien et al., 2016; Matsumoto et al., 2012). The presence of other RBDs in RGG proteins also provide them with more multivalent RNA-protein interactions and further contribute to the dynamics of RNP granules (Sanders et al., 2020).

How an RGG protein interacts with cellular RNAs and how such interactions regulate cytoplasmic mRNA translation and mRNA storage are under-investigated. Here, using an RGG-protein named Sbp1 as our model system, we studied the RGG-protein interactomes and related mRNA translation profiles at a genome-wide scale. Sbp1 is a single-stranded RNA-binding protein that contains a central RGG region flanked by two distal RNA recognition motifs (RRM) (Jong et al., 1987). Under normal physiological conditions, Sbp1 distributes throughout the cell, including to the nucleus and the cytoplasm. Our biochemical study showed that Sbp1 binds to A-rich RNA sequences (Brandariz-Nunez et al., 2018), which explains an early observation of Sbp1 co-immunoprecipating with the small nucleolar RNAs, snR10 and snR11 (Jong et al., 1987), as both snoRNAs contain A-rich RNA regions. Sbp1 co-sediments with both translating ribosomes and non-translating RNPs, and it represses general translation (Brandariz-Nunez et al., 2018). Overexpression of Sbp1 suppresses decapping defects (Segal et al., 2006). Under glucose starvation, Sbp1 accumulates in P bodies (Segal et al., 2006) and contributes to P-body dynamics (Roy et al., 2022).

In this study, using genomic and molecular approaches, we investigate how Sbp1 controls cellular mRNA translation under physiological conditions. Our results demonstrate that Sbp1 represses cytoplasmic mRNA translation and promotes mRNA storage via several distinct mechanisms including slowing ribosome movement on the mRNA, promoting polysome aggregation and stalling, and repressing translation initiation of proteins functionally important for general protein synthesis in the cell. We also show that post-translational modifications of Sbp1control the dynamics of mRNA storage and the translation. With these insights into Sbp1 function, our research establishes a framework with which the various functions of RGG-proteins can be investigated and defined.

## Results

### 1. Sbp1 slows down ribosome movement in translation elongation

To study changes of *in vivo* ribosome dynamics at a genome-wide scale and in a drug-free way, we employed the 5P-Seq methodology, which captures 5’-phosphorylated mRNA degradation intermediates, named 5P, produced by the 5’ exonuclease (Xrn1, in yeast) as it chases translating ribosomes on mRNAs (Pelechano et al., 2015). Comparing the amount of 5P intermediates captured and their corresponding positions on mRNA in the WT and ΔSbp1 cells provides valuable information on the changing ribosome dynamics associated with changes of cellular Sbp1 expression.

We obtained 5P-Seq libraries for WT and ΔSbp1 cells under normal growth conditions, when nutrients are abundant. As expected, we observe a periodic 3nt ribosomal pausing pattern as it translates consecutive codons on mRNAs. However, there is no obvious change to the preferred translation frame (Frame 1) between the 5P reads generated from the WT and ΔSbp1 cells **(Figure 1A)**. Given that the length of ribosomal footprints from the ribosomal A, P, and E-sites to its protected 5’-end are 17nt, 14nt, and 11nt, respectively, in this experiment (Pelechano et al., 2015), the peak at the -14 position with respect to the start codon AUG represents a ribosome pausing during translation initiation, when the start codon is recognized in the ribosomal P site. Likewise, the -17 peak in 5P-Seq denotes a ribosomal pause when a stop codon is encountered in ribosomal A site. As shown at the positions of -14nt in initiation and -17nt in termination (**Figure 1B)**, Sbp1 hardly changes ribosome dynamics at the start and the stop codons at the metagene level.

**Figure 1.**
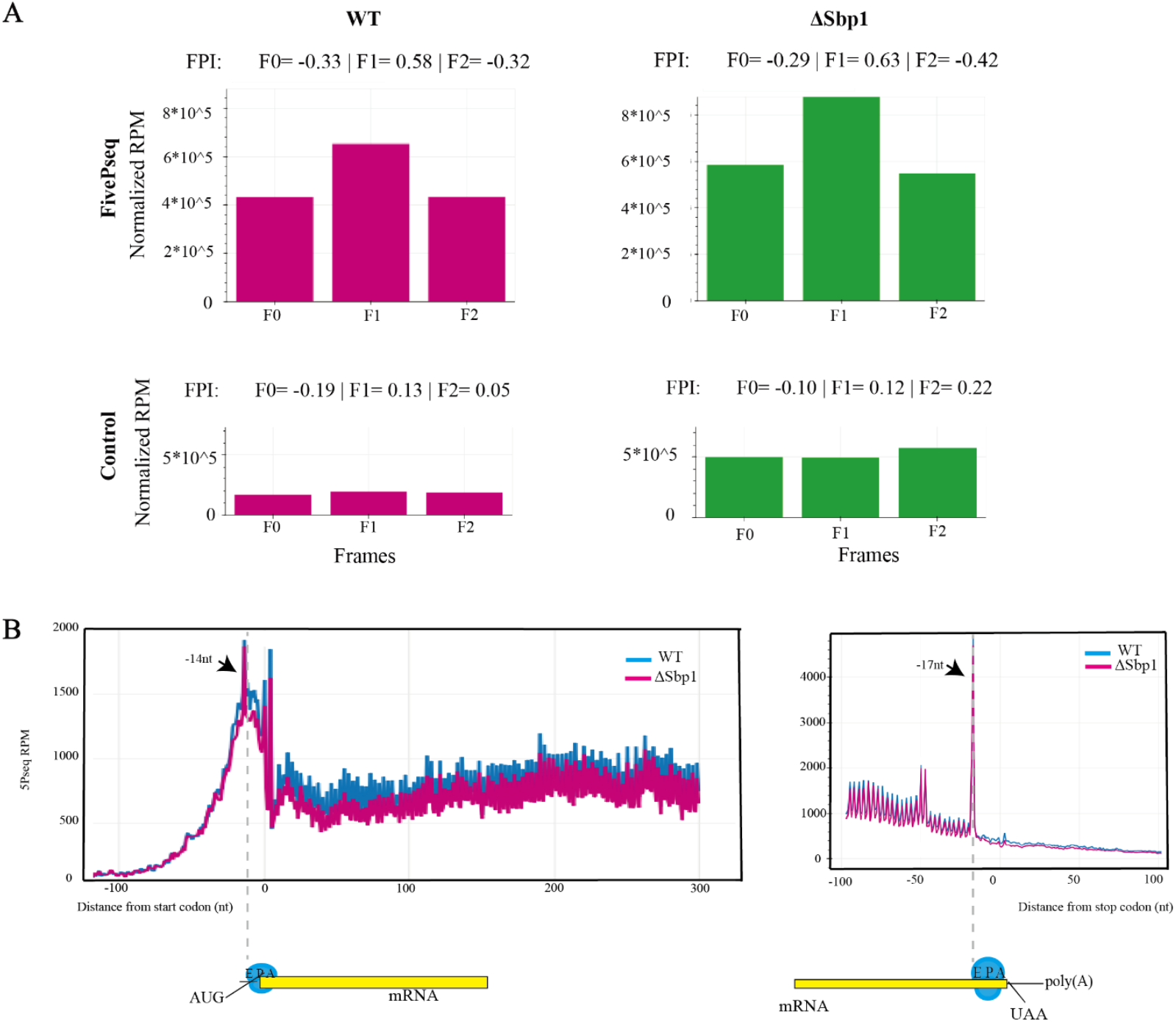
Effect of Sbp1 on cellular mRNA translation. **A The genome-wide gene frame preference in 5P-Seq**. The genome-wide gene frame preference for WT (Left) and ΔSbp1 (Right) cells in 5P-Seq under normal cell growth conditions. The top plot shows that 5P reads demonstrate F1-frame preference, which represents ribosome protected 3-nt pattern (Pelechano et al., 2015). The bottom plot is the mRNA control showing no preferred frames. The cumulative number of the 5’ end of genes in each gene frame is generated by Fivepseq (Nersisyan et al., 2020). For each figure, the horizontal axis represents frame 0,1,2 (F0, F1, F2) and the vertical axis is the normalized counts across all mRNA transcripts. Frame protection index (FPI) values and their p-value are shown at the top of the plot. FPI is computed by: FPI = log_2_(2F_i_/(F_total_- F_i_)) with i =0, 1 and 2. The FPI value larger than 0 indicates the preferred frame. **B. Absence of Sbp1 does not alter general ribosome speed in translation initiation and termination, but slows down ribosome movement in elongation under normal physiological conditions**. In 5P-Seq, no obvious accentuation of the 3nt repeating pattern was observed at the start codon (Left) and stop codon (Right) in βSbp1 cells. Changes in the height of peaks at the - 14 nt position from the start codon and -17nt position from the stop codon in WT and βSbp1 cells indicate the changes to the speed of translation initiation and termination with changes in the level of Sbp1 expression. **During translation elongation, a faster ribosome movement on mRNAs in the absence of Sbp1 is observed**. Metagene 5PSeq of wildtype and ΔSbp1 showing the 5P reads for the first 300nt from the start codon. Cumulative reads at each position were calculated and normalized to the size of the library. The peak at -14nt represents a protected mRNA region for a paused ribosome during translation initiation. Peaks shown from 0 to 300nt represent ribosome-protected regions at every 3nt during elongation. The translation dynamics of the wildtype and ΔSbp1cells are shown in blue and pink, respectively. The lower 3nt-periodicity pattern in ΔSbp1 compared to the WT indicates faster ribosome dynamics when Sbp1 protein is deleted.

In contrast, the presence of Sbp1 profoundly changes ribosome dynamics in of the elongation phase of translation. As shown in **Figure 1B (**left panel), for the first 300nt from the start codon, we observe an accentuated 3nt repeating pattern in the WT cells when Sbp1 is present compared to the one in ΔSbp1 cells. The lower peak at the -17nt position in ΔSbp1 cells indicates the faster speed of ribosome movement on mRNAs when protein Sbp1 is absent, suggesting Sbp1 represses global translation elongation. Consistent with this observation, a lower **+4** peak is shown in ΔSbp1 cells compared to that in the WT (**Figure 1B**), which represents a typical ribosome pausing when there are seven amino acids engaged in the ribosomal exit tunnel.

Furthermore, for individual amino acids involved in translation, deletion of Sbp1 resulted in an accumulation of the peak at -17nt position for Asp at the metagene level (**Figure 2**), suggesting a decreased translation speed, or even pausing when this amino acid is translated in the ribosome. Surprisingly, faster ribosomal movement was observed when Asn is located in the ribosomal P site, suggesting repressed translational pausing at this position. Clearly, the presence of Sbp1 can alter ribosome dynamics when these two amino acids are encountered.

**Figure 2.**
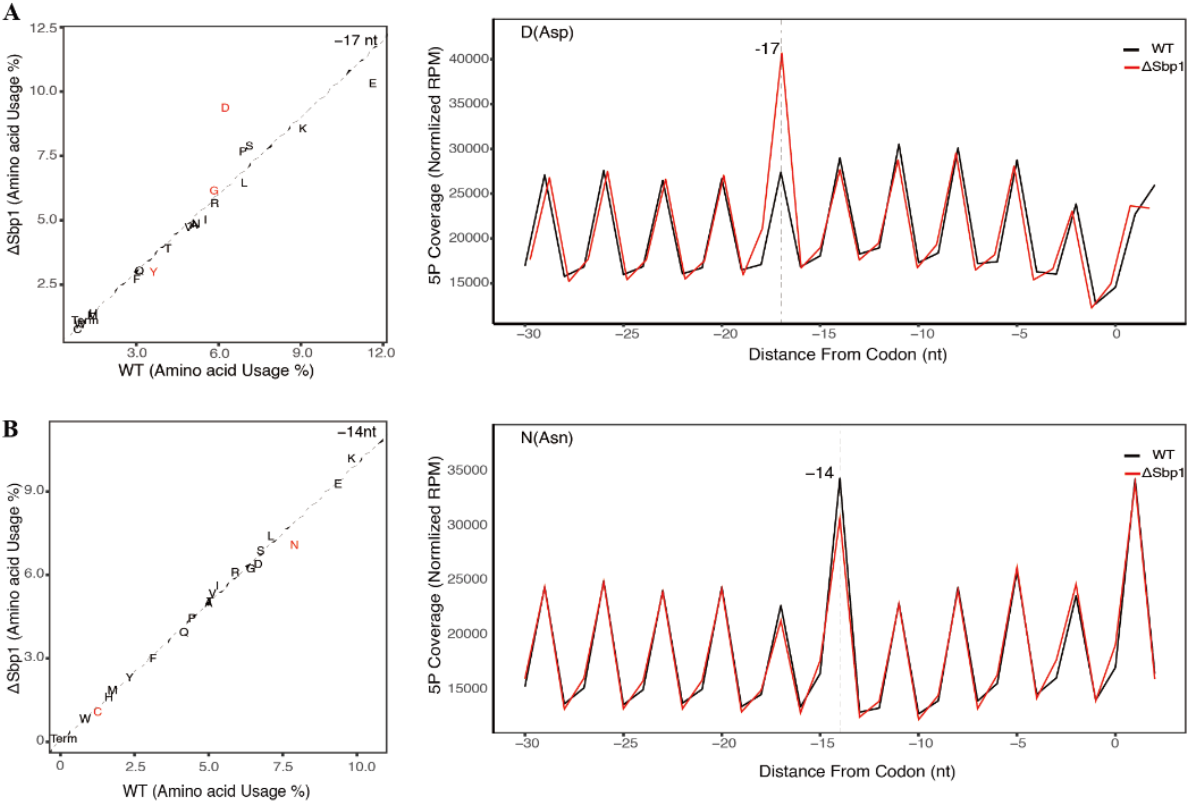
Sbp1 affects ribosome movements in an amino acid-dependent way. The differen*al ribosome pausing for all amino acids at -17 and -14 nt from the selected codon. Significantly regulated differences (adjusted P value < 0.01) are shown in red. In the absence of Sbp1, (**A**) an increased transla*on pause at Asp at the ribosomal A-site (−17nt), and (**B**) a repressed transla*on pause at Asn in the ribosomal P-site (−14nt) are observed. The metagene (- 30 to +2 window) shows the number of 5P intermediates from translation of the corresponding amino acids and their ribosomal positions in ΔSbp1 (pink lines) and WT (black lines) cells. The total number of reads was normalized to facilitate data analysis and comparison, and data were analyzed as described. The peaks at -17 nt, -14 nt, and -11 nt represent indicated codons at the ribosomal A-, P-, and E-sites, respectively. The normalized rpms for each amino acid between the WT and ΔSbp1 were compared.

### 2. Sbp1 promotes cytoplasmic polysome stalling

How does an RGG-protein, such as Sbp1, slow ribosome movement during translation? The answer may be linked to the fact that Sbp1 crosslinks with mRNA coding regions and ribosomes, as observed in our individual-nucleotide resolution UV crosslinking and immunoprecipitation (iCLIP) data (Jin lab, unpublished data). A comprehensive report of the results from the iCLIP will be reported elsewhere. Since the three domains of Sbp1 bind to RNAs synergistically and with a relatively high affinity (Brandariz-Nunez et al., 2018), we predict that tight binding of this protein to the CDS of an mRNA will impede ribosome movement during its translation.

Alternatively, it is possible that direct binding of Sbp1 to the ribosome leads to formation of a higher ordered structure, conceptually similar to colliding ribosomes (Juszkiewicz et al., 2018). To visualize the general morphology of cellular ribosomes in the presence of Sbp1, we purified Sbp1-associated ribosomal complexes and observed them after negative staining by electron microscopy. As shown in **Figure 3**, Sbp1-associated polysomes show two distinctive morphologies: an open structure, similar to beads on a string, and a closed ring-shaped structure with multiple associated ribosomes. The open beads-on-a-string structure represents actively translating ribosomes, consistent with our earlier observation that Sbp1 associates with monosomes and polysomes (Brandariz-Nunez et al., 2018). However, the closed ring-shaped and pedal-like structure is unexpected, but interesting, and we believe that it represents partially stalled polysomes due to the presence of Sbp1. This data immediately indicates that Sbp1 promotes cytoplasmic ribosome stalling.

**Figure 3.**
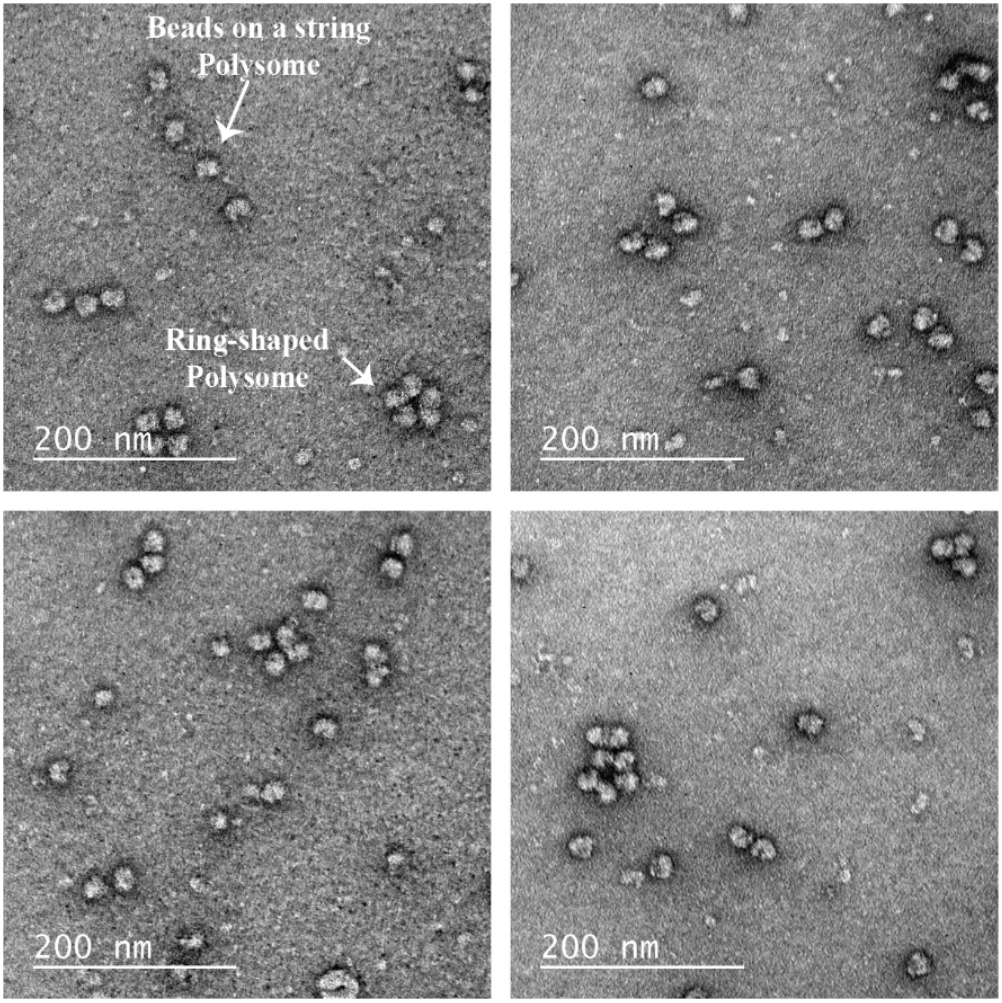
Sbp1-associated polysomes observed by electron microscopy. Typical actively translating “beads-on-a-string” polysomes and “ring-shaped” polysomes are observed in the presence of Sbp1. The open structure of “beads on a string” is an mRNA undergoing active translation, however, the closed ring-shaped structure likely represents a translation stalled mRNA.

### 3. Sbp1 represses the translation initiation of proteins functionally involved in general cellular protein production

In addition to the regulation of translation elongation as described above, translation initiation is understood to be the rate limiting step and is frequently targeted for regulation. Furthermore, data from our iCLIP show that the 5’UTRs of many cellular mRNAs are crosslinked with Sbp1 *in vivo*. Thus, the protein is likely to be functionally important for transcript-specific initiation regulation via direct mRNA association. To test this prediction, we carried out *in vitro* translation assays to determine the efficiency of translation initiation using a luciferase reporter. In this experiment, monocistronic reporter constructs were generated by cloning the native 5’UTRs of several candidate mRNA transcripts in frame with a firefly luciferase open reading frame under the control of a T7 promoter. Cap-dependent and cap-independent translation initiation, the two major pathways for the eukaryotic translation initiation, were evaluated based on the readout of luciferase reporter activities. The same amounts of RNAs were used in each translation assay, and all the measured activities were normalized to the total protein amount in the cell extract used for the assay.

Increased translation activity in the absence of Sbp1 clearly indicates an inhibitory function of this protein in both cap-dependent and cap-independent initiation of all tested RNAs (**Figure 4**). Since Sbp1 co-immunoprecipitates with snoRNAs, to rule out the possibility that the changed translation activity is due to a change in the ribosome biogenesis, we added purified Sbp1 protein into the wild-type cell extract and measured the *in vitro* translation activity in a similar manner. In the presence of increasing concentrations of purified Sbp1 ranging from 0.5µM to 4µM, gradually decreasing luciferase activity was observed (Jin Lab, unpublished data), indicating the suppressive function that this protein plays when binding directly to the 5’UTR. Consistent with this observation, our earlier investigation showed that Sbp1 inhibits *in vitro* cap-dependent and cap-independent translation initiation of the important poly(A)-binding protein in an RGG-dependent and polyA-dependent manner (Brandariz-Nunez et al., 2018).

**Figure 4.**
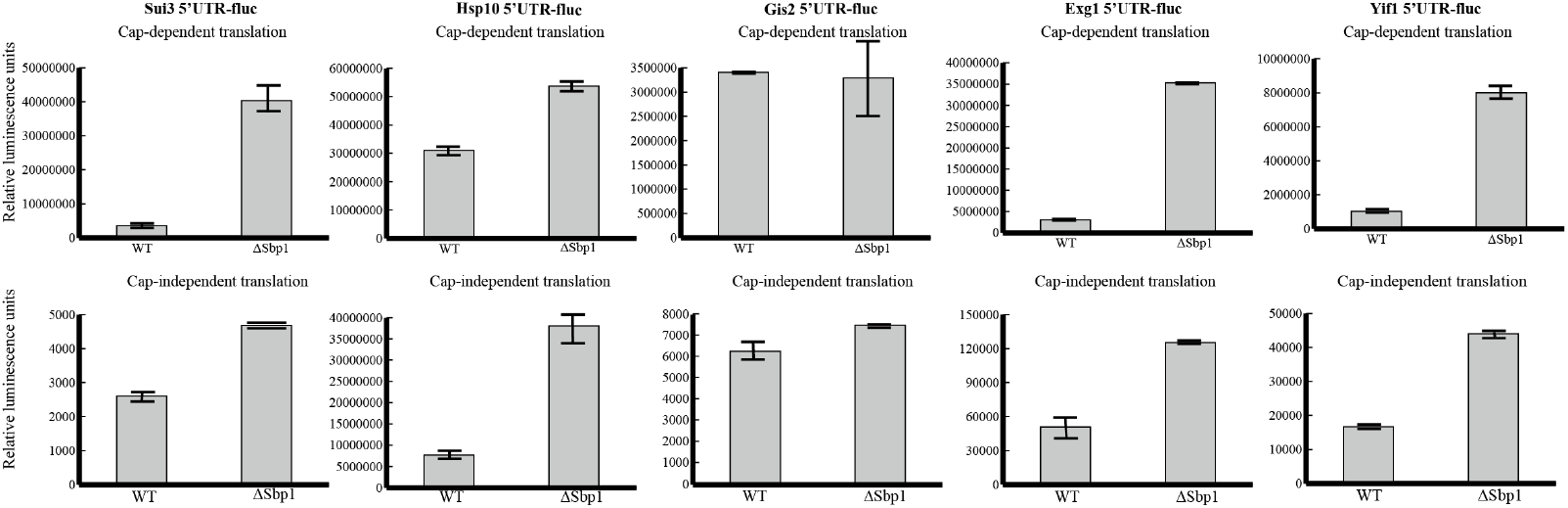
Sbp1 inhibits both cap-dependent and cap-independent translation initiation upon binding to the 5’UTR of mRNAs. The cap-dependent and cap-independent translation activities of the 5’UTRs of Hsp10, Gis2, Exg1, Sui3, and Yif1 were investigated *in vitro* using yeast cell lysates in the presence or absence of Sbp1. The mRNA reporter constructs consisted of either a capped 5′UTR or an uncapped hairpin-5’UTR upstream of the firefly luciferase gene and a poly(A) tail (5′UTR-fLuc-poly(A)). Reporter mRNAs, at a concentration of 0.66µg/µl, were incubated with either wild-type cell extracts or cell extracts lacking Sbp1, which were previously treated with nuclease to enhance the translation signal of the reporters. The vertical axis represents the relative luminescence units (RLU) of firefly luciferase, while the horizontal axis represents constructs assayed in wild-type or ΔSbp1. In all constructs, the absence of Sbp1 resulted in increased translation initiation in both the cap-dependent and cap-independent pathways. The quality of the mRNA constructs was confirmed through agarose gel electrophoresis.

It is worthwhile to point out that many of the 5’UTRs that Sbp1 directly binds to belong to mRNAs that encode important proteins necessary for efficient cellular protein synthesis. The detailed functions of the tested mRNA transcripts are summarized in **Supplemental Table 1**. For example, Sui3 is the β subunit of the translation initiation factor eIF2, which is required for start-codon recognition in translation initiation. Decreased production of the β subunit of eIF2 decreases functional eIF2 protein, which requires its three intact subunits, α, β and γ, to properly function. Conceivably, decreased availability of fully functional eIF2 proteins will result in decreased general translation that largely depends on this important initiation factor.

### 4. The methylation state of Sbp1 is important for the transition between mRNA translation and storage

Methylation at the arginine in the RGG motif fine tunes cellular functions of the RGG-proteins (Bachand, 2007; Dhar et al., 2013; Guccione and Richard, 2019; Wesche et al., 2017). To test whether post-translational modifications of Sbp1 directs the bound RNAs to translation or storage, we studied the protein contents that associate with unmethylated and methylated Sbp1, ie, Sbp1 and Sbp1^m^. In this experiment, Sbp1 was purified *in vitro* and methylation of its RGG-motif was carried out enzymatically (Brandariz-Nunez et al., 2018). These proteins were incubated with cell lysate obtained under the physiological growth conditions, and the fractions pulled-down using Sbp1 or Sbp1^m^ as a bait were analyzed by mass spectrometry. A total of 240 proteins were pulled down by Sbp1^m^, while 87 proteins were associated with unmethylated Sbp1 (**Figure 5A and Supplemental Table 2 and 3**). Gene ontology-based pathway enrichment analyses revealed that the most significantly enriched cellular components that are more closely related to or unique to the Sbp1^m^ fractions are granules, including ‘cytoplasmic stress granule’ and ‘ribonucleoprotein granule’ (adjusted p-value= 4.16E-05), as well as ‘supramolecular complex’ (adjusted p-value= 1.64E-04). In addition, the most significantly enriched biological processes in the Sbp1^m^ are all related to various metabolic processes with adjusted p-values ranging from 7.83×10 ^-11^ to 4.70×10 ^-2^. The top eight most significantly enriched classes are shown on the left panel in **Figure 5B**.

**Figure 5.**
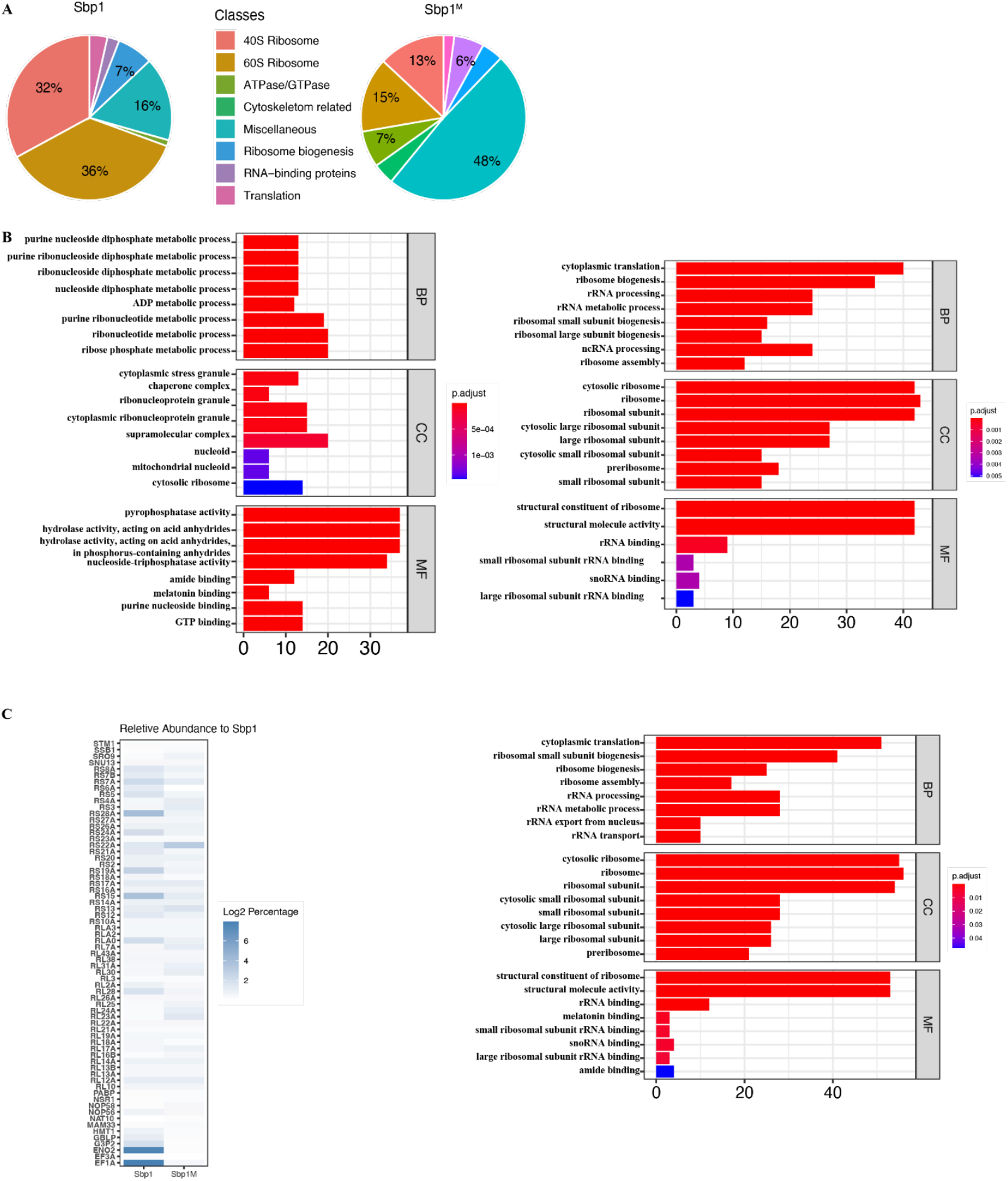
Proteins associated with Sbp1 and Sbp1^m^ that were detected mass-spectrometry. **A. The pie chart showing the main cellular component categories derived from the Gene Ontology analysis**. Pie-charts showing relative abundance of protein classes that were identified from LC-MS/MS associating with Sbp1 at different methylation states: unmethylated Sbp1 (Left) and methylated Sbp1^m^ (Right). The percentage on the chart represents proteins in the classes with high abundance (>5%). A full list of proteins that were identified and their corresponding functions are summarized in **Supplemental Table 2 and 3**. **B The gene ontology (GO) analysis for genes encoding proteins enriched in Sbp1 and Sbp1^m^ pull-down samples**. This group of genes was subjected to GO enrichment analysis, and the GO terms with Benjamini–Hochberg corrected P-value less than 0.05 considered to be significantly enriched. Biological Process (BP), Cellular Component (CC) and Molecular Function (MF) in GO sub-Ontologies are shown. p-adjusted values are shown from low to high by color transition from red to purple. The horizontal axis shows the gene ratio, which represents the percentage of genes with changes in their corresponding gene sets specified in the GO term. **C Proteins shared between Sbp1 and Sbp1^m^ pull-down samples belong to cytoplasmic translation, ribosomal components and ribosome biogenesis categories**. Normalized protein abundance in the two samples displayed as heatmap with a color spectrum ranging from white to light-blue (Left), and the GO analysis for the shared proteins in this group.

The same analyses performed on the proteins that are more related to or unique to the Sbp1-bound components highlights biological processes linked to cytoplasmic translation, ribosome biogenesis, or RNA processing. The cellular component significantly enriched in this class of proteins are all related to ribosomes, as shown on the right panel in **Figure 5B**.

It is important to note that both ribosomal components and translation factors are observed in the associated proteins pulled down by Sbp1 and Sbp1^m^, though many fewer of these proteins were found in the methylated protein Sbp1^m^ experiment (**Figure 5C**). Taken together, these data indicate an important role of RGG methylation in directing mRNAs to either translation or storage.

## Discussions

RNA-binding proteins lie at the heart of post transcriptional gene expression and regulation. RBPs with intrinsically disordered regions often interact promiscuously with other proteins and nuclear acids, which makes understanding their specific functions in different cellular pathways challenging. However, the abundance and conservation of intrinsically disordered RBPs clearly indicates their functional significance in the cell. In this study, using an abundant RNA binding protein, Sbp1, as a model system, we demonstrate how such a protein helps to establish a dynamic balance between mRNA translation and storage.

Sbp1 is a typical shuttling protein that distributes in both the nucleus and the cytosol under physiological conditions. During glucose starvation, Sbp1 accumulates in P bodies. The presence of a physiological amount of Sbp1 represses general translation in the cell and Sbp1 overexpression rescues defects in the decapping pathway (Segal et al., 2006). At the molecular level, Sbp1 possesses two well-ordered RNA recognition motifs (RRM) and an intrinsically disordered region rich in tripeptide RGG repeats. Thus, it represents a wonderful model protein to study the relationship between mRNA translation, storage and turnover.

Using the 5P-Seq genomic approach, we show that Sbp1 slows ribosome movement on mRNAs. Since Sbp1 crosslinks to the coding regions of over 20% of cellular mRNAs (Jin Lab, unpublished data), it is reasonable to assume that Sbp1 binding to mRNA coding regions alters ribosome dynamics during translation elongation. However, such a phenomenon does not change the stability of at least the majority of cellular mRNAs, because the steady-state mRNA level in the cell remains unchanged when Sbp1 is deleted (Jin Lab, unpublished data). Although it is possible that the stability of a subset of mRNA can be altered by this protein, our data clearly indicate that the main cytoplasmic function of Sbp1 lies in mRNA translation and storage.

Importantly, in the presence of Sbp1, we observed a ring-shaped polysomes when visualized by electron microscopy. Given the arrangement of this morphology, translation in the ring-shaped polysomes must slow down, if not completely stall. We predict that the ring-shaped stalled polysomes represent an intermediate state between actively translating polysomes and a translation-sequestered RNA granule. More aggregated ribosomes displaying a morula-like morphology were observed in neuronal cells previously (El Fatimy et al., 2016). Here, our data suggests that intrinsically disordered proteins promote polysome stalling under normal physiological conditions.

It is conceivable that extensive binding of Sbp1 on mRNA coding regions plays an important role in controlling the availability (and even localization) of mRNAs. In this regard, the observed ring-shaped polysomes are somewhat functionally similar to neuronal granules. Formation of such structures likely facilitates storage and transport of mRNA to specific subcellular locations where they can be translated or degraded locally.

Our data show that RBP-specific ribonucleoprotein complexes, including the observed ring-shaped polysomes, can act as a reservoir for untranslated mRNAs that can be rapidly mobilized for translation or storage by post-translational modifications of certain involved proteins, like Sbp1. Since RGG methylation alters the interactions of Sbp1 with other proteins, but leaves the Sbp1-RNA interactions largely unchanged (Brandariz-Nunez et al., 2018), it is highly likely that a different set of the proteins are recruited when Sbp1 undergoes different degrees of RGG methylation. Compared to an under-methylated Sbp1, which associates with proteins important for translation, highly methylated Sbp1 binds to a different set of proteins that are important for mRNA storage. This way, by simply targeting the post-translational modifications of an intrinsically disordered protein, cells evolved to efficiently change the gene expression program in response to internal and external stimuli.

In addition to targeting ribosomes for generalized translation repression, Sbp1 directly binds to the 5’UTRs of mRNAs and plays important roles in transcript specific translation. Sbp1 represses both cap-dependent and cap-independent translation initiation of several important proteins, such as the β subunit of the translation initiation factor eIF2 and chaperon Hasp10, necessary for efficient cellular protein synthesis. As a result, decreased production of these important proteins also contributes to the observed reduction of general translation in the cell.

It is known that Sbp1 also localizes to the nucleus, where direct association of Sbp1 with snoRNAs suggests it plays a role in ribosome biogenesis. Furthermore, in addition to methylations, Sbp1 undergoes site-specific phosphorylation at its threonine and serine residues. Since the phosphorylation of an RBP has the potential to reduce its RNA binding capacity, what are the roles of site-specific phosphorylation in this protein? Clearly, more research is required in order to appreciate the diverse cellular functions of these proteins. Dysregulation of RBP function in translation results in a variety of human diseases including metabolic diseases, muscular disorders, neurodegenerative disorders and cancer (Guo et al., 2018; Marcelo et al., 2021). Our findings and further research into the regulation of RBP functions will surely contribute to efforts to characterize related disease pathogenesis and open new therapeutic avenues.

## Acknowledgements

We thank Roy J. Carver Biotechnology Center at the University of Illinois at Urbana-Champaign for sequencing, and members in the Jin laboratory for helpful discussions. H.J. acknowledges support from the National Institute of General Medical Sciences of the NIH (R01-GM120552).

## Figures and Legends

## Methods

### Yeast strains

ΔSbp1 was constructed in the YS602 strain of *Saccharomyces cerevisiae* by homologous recombination-based replacement of Sbp1 with a kanamycin cassette using standard methods previously described (Baudin et al. 1993; Wach et al. 1994). For iCLIP, the BY4742 strain of *Saccharomyces cerevisiae* was used to chromosomally insert a 3XFLAG sequence at the C terminus of the Sbp1 coding region.

### Protein expression and purification

His-tagged or GST tagged Sbp1 was overexpressed in E. *coli* BL21 cells. Expression was induced with 0.3mM isopropyl β-D-1-thiogalactopyranoside (IPTG) with shaking at room temperature for 16hours. The proteins were initially purified with or glutathione agarose column, then further purified with ion-exchanged chromatography, next His-tagged Sbp1 was exchanged to in vitro translation buffer (22 mM Hepes-KOH, pH 7.4, 120 mM potassium acetate, 2 mM magnesium acetate, 0.75 mM ATP, 0.1 mM GTP, 25 mM creatine phosphate, 1.67 mM 1,4-dithiothreitol (DTT)) and concentrated to a final concentration of 25µM.

### In vivo methylation

Plasmids containing the GST-tagged RGG domain and type I arginine methyltransferase Hmt1 were coexpressed in E. *coli* BL21 cells. Expression was induced with 0.3mM isopropyl β- D-1-thiogalactopyranoside (IPTG) with shaking at room temperature. The methylated RGG domain was purified as described above.

### Mass spectrometry

Sbp1 proteins were expressed from *E. coli* as described above and exchanged into the buffer containing 30mM Hepes-KOH, pH 7.4, 100 mM potassium acetate, 2 mM magnesium acetate, 2mM DTT and 0.5mM PMSF. The bait protein was incubated with cell lysate obtained from cells growing under the physiological conditions for 1h at 4 °C. Methylation of the RGGs in Sbp1 was carried out enzymatically described (Brandariz-Nunez et al., 2018). After consecutive washes to remove most of the non-specific interactions, the protein complex was isolated and extracted by SDS-PAGE, then identified using liquid chromatography with tandem mass spectrometry (LC-MS/MS).

The emPAI values of each protein identified from LC-MS/MS are normalized to the total emPAI. Proteins that have 50% more normalized emPAI value between groups are considered as enriched one. For proteins identified from both Sbp1 and Sbp1^m^ groups, their abundance relative to the Sbp1 protein were measured and the results are displayed as a heatmap. The gene ontology (GO) analysis of each protein clusters were performed by R package clusterProfiler v4.0.5

### In vitro translation assays

5’UTR sequences of CHD1, BUD23 and FAS1 were identified from the BY4742 yeast strain by using the 5’RACE technique (FirstChoice RLM-RACE kit). 5’UTR sequences of Hsp10, Gis2, Exg1, Sui3 and Yif1 were obtained from the Saccharomyces genome database. A forward primer containing an XbaI cute site and reverse primers carrying BamHI cut sites were used to remove the UTRs and the first 20nt coding sequence from purified BY4742 yeast chromosomal DNA. The 5’UTR sequences were inserted between a T7 promoter and firefly luciferase gene with a short portion of the coding sequence to allow for proper folding of the 5’UTR. A 200nt polyA tail was added through PCR and *in vitro* transcription was carried out using the HiScribe T7 High Yield RNA Synthesis kit (NEB). An m7G cap was introduced by using Vaccinia capping enzyme (NEB). Cells were harvested at OD_600_ 0.5 and washed five times with lysis buffer (30mM Hepes-KOH, pH 7.4, 100 mM potassium acetate, 2 mM magnesium acetate, 2mM DTT, 0.5mM PMSF and protease inhibitor cocktail tablet (Roche))

ΔSbp1 and WT cell extracts were treated with 9,000U/ml micrococcal nuclease and 2.5mM CaCl_2_ for 7min at 26ºC (Iizuka et al., 1994). The mRNAs were unfolded at 95ºC for 30sand snap cooled prior to adding them to the translation reaction. For the in vitro translation assay using CHD1, BUD23 and FAS1 UTRs, 1µg of capped or uncapped mRNA was incubated with the treated ΔSbp1cell lysate in the presence of different concentrations of Sbp1 (0.5µM, 1µM, 2µM and 4µM). For the rest of UTRs tested, 1µg capped or uncapped mRNA were incubated with nuclease-treated ΔSbp1 and WT cell extracts. The reactions were incubated with 22 mM Hepes-KOH, pH 7.4, 120 mM potassium acetate, 2 mM magnesium acetate, 0.75 mM ATP, 0.1 mM GTP, 25 mM creatine phosphate (Invitrogen), 0.04 mM of each of the twenty amino acids, 1.67 mM 1DTT, 5 μg creatine kinase (Invitrogen) and 20U RNase inhibitor (NEB) for 1hr at 30°C. The luciferase activity was measured with ONE-Glo Luciferase reagent (Promega) with Analyst via a HT luminescence reader.

### Electron Microscopy

Samples for negative staining EM were prepared as follows: yeast cells containing chromosomally Flag tagged Sbp1 were grown to OD_600_ 0.6 at 30°C. Cells were treated with 100ug/mL cycloheximide (CHX) for 2min. Harvested cells were resuspended in 1/3^rd^ volume of lysis buffer (30mM Hepes-KOH, pH 7.4, 100 mM potassium acetate, 2 mM magnesium acetate, 2mM DTT, and protease inhibitor cocktail tablet (Roche)) and lysed in liquid nitrogen by cryogenic grinding. Cell extracts were clarified by centrifugation for 10min and loaded onto a sucrose cushion composed of 1M sucrose in lysis buffer). Ribosomes and polysomes were pelleted down by ultracentrifugation at 39k rpm for 3.5hours. Following ultracentrifugation, pellets were washed once with lysis buffer and incubated with Flag agarose beads for 1hr at 4°C. Beads were washed 5 times with lysis buffer and incubated with 100µg/ml Flag peptide to elute the Sbp1-ribosome complexes. Negative staining EM was used to observe Sbp1-ribosome complexes.

Sbp1-ribosome samples were diluted with 25mM Hepes-KOH pH 7.6, 100mM KOAc, 2.5mM Mg(OAc)2 wash buffer to an optical density of 2 A260/mL (corresponding to 40 nM of 80S) and deposited on copper grids (Electron Microscopy Sciences, CF400-CU). Grids were glow-discharged at 15 mA for 30s 4 uL of sample were incubated on the glow-discharged grids for 1min, then the grids were blotted dry, incubate with 4uL wash buffer, blot again, then negatively stained by incubation with 20uL, 20uL and 60uL of 2% Uranyl Acetate (UA) in a 3s, 3s, 30s series of consecutive incubation steps at room temperature. The excess sample was blotted away by gently touching the side of the grid with a Whatman filter paper. The grids were air-dried, then micrographs were acquired on a JEOL2100 transmission electron microscope (Thermo Ficher Scientific) operated at 120 kV using a room temperature side entry holder. Images were collected with 2k x 2k pixels Gatan UltraScan CCD camera at a nominal magnification of 40kx

### 5P-Seq

ΔSbp1 and wildtype cellswere grown in YPD to OD_600_ 0.5, harvested, then the total cellular RNA was extracted by the Trizol method. More than 50ug of total RNA was used to generate standard 5PSeq libraries according to Pelechano et al., 2016. Total RNA extracts were treated with 5U PNK and ligated with 10µM RNA oligo linkers at 16°C overnight. Poly(A) mRNA was enriched using 100µl oligo(dT) (Dynabeads oligo (dT)_25_) for 5P samples while randomly generated fragments of total RNA were prepared as controls. Both 5P and control RNAs were fragmented at 80°C for 5min. Following reverse transcription and second strand generation, cDNA libraries were incubated with M-280 streptavidin beads overnight at 4 ºC, instead of 30min at room temperature for maximal binding of the cDNA. The final DNA adapter ligation reaction was carried out for 2hrs. Enriched fragments between 300-500nt were sequenced through single end high throughput sequencing with Illumina NovaSeq 6000 system.

### 5P-Seq data analysis

Adapters were trimmed from sequenced reads using cutadapt v1.18. Then unique molecular identifiers (UMI) were extracted and deduplicated using UMI tools v1.01. Reads were aligned to the Saccharomyces cerevisiae. R64-1-1.108 genome (Ensemble) by STAR v2.7.3a. BedGraph profiles and RNA content were separated by bedtools v2.29.2 and samtools v1.10.2. mRNA potion of samples analysis by fivepseq package v1.0.0. The following analysis and plot making was conducted R v4.1.0 with package ggplot2 v3.4.2 and clusterProfiler v4.0.5

