## Supplemental Materials for "An intrinsically Disordered RNA Binding Protein Modulates mRNA Translation and Storage"

### Supplemental Information

**Supplemental Table 1. Main cellular functions for proteins encoding candidate mRNAs under the biochemical investigation.**

| Gene names | Protein names | Functions |
|---|---|---|
| Sui3 | eIF2-β subunit | Sui3 is the beta subunit of the translation initiation factor eIF2 which is important for the start codon recognition (Donahue, Cigan, Pabich, & Valavicius, 1988; Laurino, Thompson, Pacheco, & Castilho, 1999). It is conceivable the decreased production of the beta subunit of eIF2 where lead to a decreased amount of functional eIF2 protein, which result in an inhibition of the general translation that largely depends on these important initiation factor (Hinnebusch & Lorsch, 2012). |
| Hsp10 | Hsp10 | Hsp10 is a mitochondrial chaperonin, involved in protein folding and sorting in the mitochondria (Böttinger et al., 2015; Gomez-Llorente et al., 2020; Höhfeld & Hartl, 1994). |
| Gis2 | Gis2 | Gis2 is a protein associates with polysomes and play important functions in translation regulation under stress conditions (Rojas et al., 2012; Scherrer, Femmer, Schiess, Aebersold, & Gerber, 2011). |
| Exg1 | Exg1 | Exg1 is a major glucanase of the cell wall, involved in yeast cell wall beta-glucan assembly (Nebreda, Villa, Villanueva, & del Rey, 1986). |
| Yif1 | Yif1 | Yif1 is an integral transmembrane protein, important for the fusion of ER-derived COPII transport vehicles with the Golgi (Barrowman, Wang, Zhang, & Ferro-Novick, 2003; Heidtman, Chen, Collins, & Barlowe, 2005). |

**Supplemental Table 2. Proteins associated with methylated Sbp1 identified by mass spectrometry.**

| Gene names | Protein names | Types | Descriptions |
|---|---|---|---|
| SBP1 | SSBP1 | Granule, P bodies/Stress Granules | Single-stranded nucleic acid-binding protein |
| TEF1 | EF1A | Cytoskeleton | Elongation factor 1-alpha |
| VMA2 | VATB | ATPase | V-type proton ATPase subunit B |
| ENO2 | ENO2 | Miscellaneous | Enolase 2 |
| RPS5 | RS5 | Ribosome | 40S ribosomal protein S5 |
| TDH3 | G3P3 | Miscellaneous | Glyceraldehyde-3-phosphate dehydrogenase 3 |
| TDH2 | G3P2 | Miscellaneous | Glyceraldehyde-3-phosphate dehydrogenase 2 |
| ATP1 | ATPA | ATPase | ATP synthase subunit alpha, mitochondrial |
| CPR1 | CYPH | Miscellaneous | Peptidyl-prolyl cis-trans isomerase |
| ATP2 | ATPB | ATPase | ATP synthase subunit beta, mitochondrial |
| FBA1 | ALF | Miscellaneous | Fructose-bisphosphate aldolase |
| PGK1 | PGK | Miscellaneous | Phosphoglycerate kinase |
| KAP123 | IMB4 | Granule, Ribonucleoprotein granule | Importin subunit beta-4 |
| YKT6 | YKT6 | Miscellaneous | Synaptobrevin homolog YKT6 |
| RPS19A | RS19A | Ribosome | 40S ribosomal protein S19-A |
| RPS19B | RS19B | Ribosome | 40S ribosomal protein S19-B |
| PDC1 | PDC1 | Miscellaneous | Pyruvate decarboxylase isozyme 1 |
| NOP56 | NOP56 | Ribosome | Nucleolar protein 56 |
| CCT4 | TCPD | Miscellaneous | T-complex protein 1 subunit delta |
| HSP60 | HSP60 | Miscellaneous | Heat shock protein 60, mitochondrial |
| ASC1 | GBLP | Ribosome | Guanine nucleotide-binding protein subunit beta-like protein |
| TUB2 | TBB | Cytoskeleton | Tubulin beta chain |
| DED1 | DED1 | Stress granule | ATP-dependent RNA helicase DED1 |
| RPS4A | RS4A | Ribosome | 40S ribosomal protein S4-A |
| IDH2 | IDH2 | Miscellaneous | Isocitrate dehydrogenase [NAD] subunit 2, mitochondrial |
| RPS15 | RS15 | Ribosome | 40S ribosomal protein S15 |
| RPP0 | RLA0 | Ribosome | 60S acidic ribosomal protein P0 |
| RPL4B | RL4B | Ribosome | 60S ribosomal protein L4-B |
| RPL12A | RL12A | Ribosome | 60S ribosomal protein L12-A |
| TUB3 | TBA3 | Cytoskeleton | Tubulin alpha-3 chain |
| TUB1 | TBA1 | Cytoskeleton | Tubulin alpha-1 chain |
| LSB3 | LSB3 | Cytoskeleton | LAS seventeen-binding protein 3 |
| RPS12 | RS12 | Ribosome | 40S ribosomal protein S12 |
| RPS2 | RS2 | Ribosome | 40S ribosomal protein S2 |
| TCP1 | TCPA | Miscellaneous | T-complex protein 1 subunit alpha |
| SAM1 | METK1 | Stress granule | S-adenosylmethionine synthase 1 |

| | | | |
|---|---|---|---|
| CDC19 | KPYK1 | Miscellaneous | Pyruvate kinase 1 |
| RPS31 | RS31 | Ribosome | Ubiquitin-40S ribosomal protein S31 |
| GPM1 | PMG1 | Miscellaneous | Phosphoglycerate mutase 1 |
| RPL8B | RL8B | Ribosome | 60S ribosomal protein L8-B |
| CCT3 | TCPG | Miscellaneous | T-complex protein 1 subunit gamma |
| RPL14A | RL14A | Ribosome | 60S ribosomal protein L14-A |
| RPS21B | RS21B | Ribosome | 40S ribosomal protein S21-B |
| RPS21A | RS21A | Ribosome | 40S ribosomal protein S21-A |
| SSA2 | HSP72 | Ribosome | Heat shock protein SSA2 |
| RPS17A | RS17A | Ribosome | 40S ribosomal protein S17-A |
| SRO9 | SRO9 | Granule, P bodies | RNA-binding protein SRO9 |
| SHM2 | GLYC | Miscellaneous | Serine hydroxymethyltransferase, cytosolic |
| RPS7B | RS7B | Ribosome | 40S ribosomal protein S7-B |
| RPS7A | RS7A | Ribosome | 40S ribosomal protein S7-A |
| VPS1 | VPS1 | Cytoskeleton | Vacuolar protein sorting-associated protein 1 |
| RPS1B | RS3A2 | Ribosome | 40S ribosomal protein S1-B |
| RPS1A | RS3A1 | Ribosome | 40S ribosomal protein S1-A |
| RPL17A | RL17A | Ribosome | 60S ribosomal protein L17-A |
| YPK1 | YPK1 | Miscellaneous | Serine/threonine-protein kinase YPK1 |
| EFT1 | EF2 | Miscellaneous | Elongation factor 2 |
| URA7 | URA7 | Miscellaneous | CTP synthase 1 |
| RPS8A | RS8A | Ribosome | 40S ribosomal protein S8-A |
| VMA1 | VATA | ATPase | V-type proton ATPase catalytic subunit A |
| STM1 | STM1 | Ribosome | Suppressor protein STM1 |
| RPL19A | RL19A | Ribosome | 60S ribosomal protein L19-A |
| VMA8 | VATD | ATPase | V-type proton ATPase subunit D |
| GET3 | GET3 | Miscellaneous | ATPase GET3 |
| RPS22A | RS22A | Ribosome | 40S ribosomal protein S22-A |
| YEF3 | EF3A | Granule, Ribonucleoprotein granule | Elongation factor 3A |
| RPS6A | RS6A | Ribosome | 40S ribosomal protein S6-A |
| RPS24A | RS24A | Ribosome | 40S ribosomal protein S24-A |
| RRP12 | RRP12 | Ribosome | Ribosomal RNA-processing protein 12 |
| GGA2 | GGA2 | Miscellaneous | ADP-ribosylation factor-binding protein GGA2 |
| RPL43A | RL43A | Ribosome | 60S ribosomal protein L43-A |
| RPL25 | RL25 | Ribosome | 60S ribosomal protein L25 |
| NOG2 | NOG2 | Ribosome | Nucleolar GTP-binding protein 2 |
| CCT6 | TCPZ | Miscellaneous | T-complex protein 1 subunit zeta |
| HMT1 | HMT1 | Miscellaneous | Protein arginine N-methyltransferase 1 |
| RPL22A | RL22A | Ribosome | 60S ribosomal protein L22-A |
| CCT8 | TCPQ | Miscellaneous | T-complex protein 1 subunit theta |
| RPS13 | RS13 | Ribosome | 40S ribosomal protein S13 |
| RVS167 | RV167 | Cytoskeleton | Reduced viability upon starvation protein 167 |

| | | | |
|---|---|---|---|
| DHH1 | DHH1 | Granule, P bodies | ATP-dependent RNA helicase DHH1 |
| PAB1 | PABP | Stress granue | Polyadenylate-binding protein, cytoplasmic and nuclear |
| RPS16A | RS16A | Ribosome | 40S ribosomal protein S16-A |
| RPS3 | RS3 | Ribosome | 40S ribosomal protein S3 |
| RPL10 | RL10 | Ribosome | 60S ribosomal protein L10 |
| RPS26A | RS26A | Ribosome | 40S ribosomal protein S26-A |
| RPL13B | RL13B | Ribosome | 60S ribosomal protein L13-B |
| RPL13A | RL13A | Ribosome | 60S ribosomal protein L13-A |
| NPL3 | NOP3 | Granule, Stress granule | Nucleolar protein 3 |
| ADH1 | ADH1 | Miscellaneous | Alcohol dehydrogenase 1 |
| RPS0A | RSSA1 | Ribosome | 40S ribosomal protein S0-A |
| RPS20 | RS20 | Ribosome | 40S ribosomal protein S20 |
| IDH1 | IDH1 | Miscellaneous | Isocitrate dehydrogenase [NAD] subunit 1, mitochondrial |
| RPL2A | RL2A | Ribosome | 60S ribosomal protein L2-A |
| RPP1B | RLA3 | Ribosome | 60S acidic ribosomal protein P1-beta |
| RPL16B | RL16B | Ribosome | 60S ribosomal protein L16-B |
| RPL1A | RL1A | Ribosome | 60S ribosomal protein L1-A |
| NOP58 | NOP58 | Ribosome | Nucleolar protein 58 |
| COG8 | COG8 | Miscellaneous | Conserved oligomeric Golgi complex subunit 8 |
| RPL3 | RL3 | Ribosome | 60S ribosomal protein L3 |
| USO1 | USO1 | Cytoskeleton | Intracellular protein transport protein USO1 |
| ILV2 | ILVB | Miscellaneous | Acetolactate synthase catalytic subunit, mitochondrial |
| RPL28 | RL28 | Ribosome | 60S ribosomal protein L28 |
| RNR1 | RIR1 | Miscellaneous | Ribonucleoside-diphosphate reductase large chain 1 |
| RPL5 | RL5 | Ribosome | 60S ribosomal protein L5 |
| PRE8 | PSA2 | Miscellaneous | Proteasome subunit alpha type-2 |
| TY1B | TY1AB | Miscellaneous | Transposon TyH3 Gag-Pol polyprotein |
| RPS23A | RS23A | Ribosome | 40S ribosomal protein S23-A |
| ARF1 | ARF1 | Miscellaneous | ADP-ribosylation factor 1 |
| ARF2 | ARF2 | Miscellaneous | ADP-ribosylation factor 2 |
| NUG1 | NUG1 | Ribosome | Nuclear GTP-binding protein NUG1 |
| RPS28A | RS28A | Ribosome | 40S ribosomal protein S28-A |
| GPP1 | GPP1 | Miscellaneous | Glycerol-1-phosphate phosphohydrolase 1 |
| ACB1 | ACBP | Miscellaneous | Acyl-CoA-binding protein |
| LYS12 | LYS12 | Miscellaneous | Homoisocitrate dehydrogenase, mitochondrial |
| PSA1 | MPG1 | Miscellaneous | Mannose-1-phosphate guanyltransferase |
| COQ1 | COQ1 | Miscellaneous | Hexaprenyl pyrophosphate synthase, mitochondrial |
| TIF1 | IF4A | Granule | ATP-dependent RNA helicase eIF4A |

| | | | |
|---|---|---|---|
| HEM1 | HEM1 | Miscellaneous | 5-aminolevulinate synthase, mitochondrial |
| GUK1 | KGUA | Miscellaneous | Guanylate kinase |
| SCY1 | SCY1 | Miscellaneous | Protein kinase-like protein SCY1 |
| ADE17 | PUR92 | Miscellaneous | Bifunctional purine biosynthesis protein ADE17 |
| PMA1 | PMA1 | Miscellaneous | Plasma membrane ATPase 1 |
| SNU13 | SNU13 | Ribosome | 13 kDa ribonucleoprotein-associated protein |
| RPO21 | RPB1 | Ribonucleoprotein granule | DNA-directed RNA polymerase II subunit RPB1 |
| YPT7 | YPT7 | Miscellaneous | GTP-binding protein YPT7 |
| ILV1 | THDH | Miscellaneous | Threonine dehydratase, mitochondrial |
| BEM1 | BEM1 | Cytoskeleton | Bud emergence protein 1 |
| PUF3 | PUF3 | Miscellaneous | mRNA-binding protein PUF3 |
| SPA2 | SPA2 | Cytoskeleton | Protein SPA2 |
| GFA1 | GFA1 | Miscellaneous | Glutamine--fructose-6-phosphate aminotransferase [isomerizing] |
| RPS14A | RS14A | Ribosome | 40S ribosomal protein S14-A |
| PDA1 | ODPA | Miscellaneous | Pyruvate dehydrogenase E1 component subunit alpha, mitochondrial |
| ACT1 | ACT | Cytoskeleton | Actin |
| CHD1 | CHD1 | Miscellaneous | Chromo domain-containing protein 1 |
| CDC39 | NOT1 | Granule, P bodies | General negative regulator of transcription subunit 1 |
| SDA1 | SDA1 | Miscellaneous | Protein SDA1 |
| DNM1 | DNM1 | Miscellaneous | Dynamin-related protein DNM1 |
| GRX1 | GLRX1 | Miscellaneous | Glutaredoxin-1 |
| MRP4 | RT04 | Ribosome | 37S ribosomal protein MRP4, mitochondrial |
| HSC82 | HSC82 | Miscellaneous | ATP-dependent molecular chaperone HSC82 |
| PUF2 | PUF2 | Miscellaneous | mRNA-binding protein PUF2 |
| TRP5 | TRP | Miscellaneous | Tryptophan synthase |
| YHR020W | YHI0 | Miscellaneous | Putative proline--tRNA ligase YHR020W |
| CPR3 | CYPC | Miscellaneous | Peptidyl-prolyl cis-trans isomerase C, mitochondrial |
| ADK1 | KAD2 | Miscellaneous | Adenylate kinase |
| RPL21A | RL21A | Ribosome | 60S ribosomal protein L21-A |
| DBP5 | DBP5 | Granule, Ribonucleoprotein granule | ATP-dependent RNA helicase DBP5 |
| NAP1 | NAP1 | Cytoskeleton | Nucleosome assembly protein |
| SER1 | SERC | Miscellaneous | Phosphoserine aminotransferase |
| JSN1 | JSN1 | Miscellaneous | Protein JSN1 |
| CDC9 | DNLI1 | Miscellaneous | DNA ligase 1 |
| GRX2 | GLRX2 | Miscellaneous | Glutaredoxin-2, mitochondrial |
| CCT5 | TCPE | Miscellaneous | T-complex protein 1 subunit epsilon |
| COP1 | COPA | Miscellaneous | Coatomer subunit alpha |

| | | | |
|---|---|---|---|
| MOT2 | NOT4 | Miscellaneous | General negative regulator of transcription subunit 4 |
| RPA49 | RPA49 | Miscellaneous | DNA-directed RNA polymerase I subunit RPA49 |
| YSC84 | YSC84 | Cytoskeleton | Protein YSC84 |
| NOG1 | NOG1 | Ribosome | Nucleolar GTP-binding protein 1 |
| RPL18A | RL18A | Ribosome | 60S ribosomal protein L18-A |
| RPS27A | RS27A | Ribosome | 40S ribosomal protein S27-A |
| RPL30 | RL30 | Ribosome | 60S ribosomal protein L30 |
| YPT52 | YPT52 | Miscellaneous | GTP-binding protein YPT52 |
| RPL38 | RL38 | Ribosome | 60S ribosomal protein L38 |
| ALD6 | ALDH6 | Miscellaneous | Magnesium-activated aldehyde dehydrogenase, cytosolic |
| RPL24A | RL24A | Ribosome | 60S ribosomal protein L24-A |
| PFY1 | PROF | Cytoskeleton | Profilin |
| SSC1 | HSP77 | Miscellaneous | Heat shock protein SSC1, mitochondrial |
| FAA4 | LCF4 | Granule | Long-chain-fatty-acid--CoA ligase 4 |
| NOP14 | NOP14 | Ribosome | Nucleolar complex protein 14 |
| SES1 | SYSC | Ribonucleoprote in granule | Serine--tRNA ligase, cytoplasmic |
| CLU1 | CLU | Ribonucleoprote in granule | Clustered mitochondria protein 1 |
| RPP2A | RLA2 | Ribosome | 60S acidic ribosomal protein P2-alpha |
| YRA1 | YRA1 | Miscellaneous | RNA annealing protein YRA1 |
| CYS3 | CYS3 | Miscellaneous | Cystathionine gamma-lyase |
| SEC65 | SEC65 | Miscellaneous | Signal recognition particle subunit SEC65 |
| NAB6 | NAB6 | Granule, Stress granule | RNA-binding protein NAB6 |
| RPL31A | RL31A | Ribosome | 60S ribosomal protein L31-A |
| CSR1 | CSR1 | Miscellaneous | Phosphatidylinositol transfer protein CSR1 |
| RPL33A | RL33A | Ribosome | 60S ribosomal protein L33-A |
| RPS18A | RS18A | Ribosome | 40S ribosomal protein S18-A |
| NOC2 | NOC2 | Ribosome | Nucleolar complex protein 2 |
| SGD1 | SGD1 | Miscellaneous | Suppressor of glycerol defect protein 1 |
| RPC10 | RPAB4 | Miscellaneous | DNA-directed RNA polymerases I, II, and III subunit RPABC4 |
| EFR3 | EFR3 | Miscellaneous | Protein EFR3 |
| SEC16 | SEC16 | Miscellaneous | COPII coat assembly protein SEC16 |
| PFK2 | PFKA2 | Miscellaneous | ATP-dependent 6-phosphofructokinase subunit beta |
| ATP3 | ATPG | ATPase | ATP synthase subunit gamma, mitochondrial |
| MAM33 | MAM33 | Miscellaneous | Mitochondrial acidic protein MAM33 |
| RPL11A | RL11A | Ribosome | 60S ribosomal protein L11-A |
| GSP1 | GSP1 | Miscellaneous | GTP-binding nuclear protein GSP1/CNR1 |
| TPI1 | TPIS | Miscellaneous | Triosephosphate isomerase |
| PUF4 | PUF4 | Miscellaneous | Pumilio homology domain family member 4 |

| | | | |
|---|---|---|---|
| VMA5 | VATC | ATPase | V-type proton ATPase subunit C |
| URA2 | PYR1 | Miscellaneous | Protein URA2 |
| KRE33 | NAT10 | Ribosome | RNA cytidine acetyltransferase |
| RNR4 | RIR4 | Miscellaneous | Ribonucleoside-diphosphate reductase small chain 2 |
| BRE5 | BRE5 | Granule, Stress granule | UBP3-associated protein BRE5 |
| ARP3 | ARP3 | Cytoskeleton | Actin-related protein 3 |
| PPN1 | PPN1 | Miscellaneous | Endopolyphosphatase |
| RPL15A | RL15A | Ribosome | 60S ribosomal protein L15-A |
| BMH1 | BMH1 | Granule | Protein BMH1 |
| SCY_3392 | YKR18 | Miscellaneous | Mitochondrial outer membrane protein SCY_3392 |
| HYP2 | IF5A1 | Miscellaneous | Eukaryotic translation initiation factor 5A-1 |
| RPL23A | RL23A | Ribosome | 60S ribosomal protein L23-A |
| YNL208W | YNU8 | Miscellaneous | Uncharacterized protein YNL208W |
| RPN7 | RPN7 | Miscellaneous | 26S proteasome regulatory subunit RPN7 |
| RPB10 | RPAB5 | Miscellaneous | DNA-directed RNA polymerases I, II, and III subunit RPABC5 |
| RPL36A | RL36A | Ribosome | 60S ribosomal protein L36-A |
| MCM4 | MCM4 | Miscellaneous | DNA replication licensing factor MCM4 |
| RPS10A | RS10A | Ribosome | 40S ribosomal protein S10-A |
| IMD1 | IMDH1 | Miscellaneous | Putative inosine-5'-monophosphate dehydrogenase 1 |
| TBF1 | TBF1 | Miscellaneous | Protein TBF1 |
| GAR1 | GAR1 | Miscellaneous | H/ACA ribonucleoprotein complex subunit 1 |
| RPL7A | RL7A | Ribosome | 60S ribosomal protein L7-A |
| PPH22 | PP2A2 | Miscellaneous | Serine/threonine-protein phosphatase PP2A-2 catalytic subunit |
| STU1 | STU1 | Cytoskeleton | Protein STU1 |
| SSB1 | SSB1 | Ribosome | Ribosome-associated molecular chaperone SSB1 |
| RPL35A | RL35A | Ribosome | 60S ribosomal protein L35-A |
| MAK21 | MAK21 | Ribosome | Ribosome biogenesis protein MAK21 |
| RRP3 | RRP3 | Miscellaneous | ATP-dependent rRNA helicase RRP3 |
| NSR1 | NSR1 | Miscellaneous | Nuclear localization sequence-binding protein |
| IPP1 | IPYR | Miscellaneous | Inorganic pyrophosphatase |
| ARP2 | ARP2 | Cytoskeleton | Actin-related protein 2 |
| FPR2 | FKBP2 | Miscellaneous | Peptidyl-prolyl cis-trans isomerase FPR2 |
| GCV3 | GCSH | Miscellaneous | Glycine cleavage system H protein, mitochondrial |
| RET3 | COPZ | Miscellaneous | Coatomer subunit zeta |
| SSE1 | HSP7F | Ribosome | Heat shock protein homolog SSE1 |
| RPT2 | PRS4 | Miscellaneous | 26S proteasome regulatory subunit 4 homolog |
| CDC3 | CDC3 | Cytoskeleton | Cell division control protein 3 |

| | | | |
|---|---|---|---|
| SCH9 | SCH9 | Miscellaneous | Serine/threonine-protein kinase SCH9 |
| PDB1 | ODPB | Miscellaneous | Pyruvate dehydrogenase E1 component subunit beta, mitochondrial |
| STE5 | STE5 | Miscellaneous | Protein STE5 |
| DBP9 | DBP9 | Miscellaneous | ATP-dependent RNA helicase DBP9 |
| RPL26A | RL26A | Ribosome | 60S ribosomal protein L26-A |
| RDL2 | RDL2 | Miscellaneous | Thiosulfate sulfurtransferase RDL2, mitochondrial |
| MDN1 | MDN1 | Ribosome | Midasin |
| PRE2 | PSB5 | Miscellaneous | Proteasome subunit beta type-5 |
| YML6 | RL4P | Ribosome | 54S ribosomal protein YmL6, mitochondrial |
| EDE1 | EDE1 | Cytoskeleton | EH domain-containing and endocytosis protein 1 |
| YPT1 | YPT1 | Miscellaneous | GTP-binding protein YPT1 |
| GUA1 | GUAA | Miscellaneous | GMP synthase [glutamine-hydrolyzing] |

**Supplemental Table 3. Proteins associated with unmethylated Sbp1 identified by mass spectrometry.**

| Gene names | Protein names | Types | Descriptions |
|---|---|---|---|
| SBP1 | SSBP1 | Granule, P bodies and stress granules | Single-stranded nucleic acid-binding protein |
| RPS22A | RS22A | Ribosome | 40S ribosomal protein S22-A |
| RPS13 | RS13 | Ribosome | 40S ribosomal protein S13 |
| RPL23A | RL23A | Ribosome | 60S ribosomal protein L23-A |
| RPS3 | RS3 | Ribosome | 40S ribosomal protein S3 |
| RPS17A | RS17A | Ribosome | 40S ribosomal protein S17-A |
| RPS15 | RS15 | Ribosome | 40S ribosomal protein S15 |
| RPS7A | RS7A | Ribosome | 40S ribosomal protein S7-A |
| RPL7A | RL7A | Ribosome | 60S ribosomal protein L7-A |
| RPL4A | RL4A | Ribosome | 60S ribosomal protein L4-A |
| RPL30 | RL30 | Ribosome | 60S ribosomal protein L30 |
| RPS4A | RS4A | Ribosome | 40S ribosomal protein S4-A |
| RPL24A | RL24A | Ribosome | 60S ribosomal protein L24-A |
| RPL12A | RL12A | Ribosome | 60S ribosomal protein L12-A |
| RPL33B | RL33B | Ribosome | 60S ribosomal protein L33-B |
| RPL31A | RL31A | Ribosome | 60S ribosomal protein L31-A |
| RPP0 | RLA0 | Ribosome | 60S acidic ribosomal protein P0 |
| RPS6A | RS6A | Ribosome | 40S ribosomal protein S6-A |
| RPL17A | RL17A | Ribosome | 60S ribosomal protein L17-A |
| RPS26A | RS26A | Ribosome | 40S ribosomal protein S26-A |
| IMD3 | IMDH3 | Miscellaneous | Inosine-5'-monophosphate dehydrogenase 3 |
| RPS20 | RS20 | Ribosome | 40S ribosomal protein S20 |
| RPS14A | RS14A | Ribosome | 40S ribosomal protein S14-A |
| RPS8A | RS8A | Ribosome | 40S ribosomal protein S8-A |
| RPL14A | RL14A | Ribosome | 60S ribosomal protein L14-A |
| RPS24A | RS24A | Ribosome | 40S ribosomal protein S24-A |
| RPS12 | RS12 | Ribosome | 40S ribosomal protein S12 |
| RPS28A | RS28A | Ribosome | 40S ribosomal protein S28-A |
| SRO9 | SRO9 | Granule | RNA-binding protein SRO9 |
| RPS19A | RS19A | Ribosome | 40S ribosomal protein S19-A |
| RPL25 | RL25 | Ribosome | 60S ribosomal protein L25 |
| RPS16A | RS16A | Ribosome | 40S ribosomal protein S16-A |
| RPS5 | RS5 | Ribosome | 40S ribosomal protein S5 |
| RPL10 | RL10 | Ribosome | 60S ribosomal protein L10 |
| RPL28 | RL28 | Ribosome | 60S ribosomal protein L28 |
| RPL38 | RL38 | Ribosome | 60S ribosomal protein L38 |
| RPS2 | RS2 | Ribosome | 40S ribosomal protein S2 |
| RPS27A | RS27A | Ribosome | 40S ribosomal protein S27-A |
| RPS21A | RS21A | Ribosome | 40S ribosomal protein S21-A |
| RPL20A | RL20A | Ribosome | 60S ribosomal protein L20-A |
| RPL43A | RL43A | Ribosome | 60S ribosomal protein L43-A |
| RPS7B | RS7B | Ribosome | 40S ribosomal protein S7-B |

| | | | |
|---|---|---|---|
| RPL19A | RL19A | Ribosome | 60S ribosomal protein L19-A |
| RPP1B | RLA3 | Ribosome | 60S acidic ribosomal protein P1-beta |
| RPL16B | RL16B | Ribosome | 60S ribosomal protein L16-B |
| RPP2A | RLA2 | Ribosome | 60S acidic ribosomal protein P2-alpha |
| IMD4 | IMDH4 | Miscellaneous | Inosine-5'-monophosphate dehydrogenase 4 |
| RPL13B | RL13B | Ribosome | 60S ribosomal protein L13-B |
| RPL13A | RL13A | Ribosome | 60S ribosomal protein L13-A |
| TEF1 | EF1A | Miscellaneous | Elongation factor 1-alpha |
| RPS10A | RS10A | Ribosome | 40S ribosomal protein S10-A |
| SNU13 | SNU13 | Ribosome | 13 kDa ribonucleoprotein-associated protein |
| RPL2A | RL2A | Ribosome | 60S ribosomal protein L2-A |
| NOP56 | NOP56 | Ribosome | Nucleolar protein 56 |
| RPS0A | RSSA1 | Ribosome | 40S ribosomal protein S0-A |
| RPL22A | RL22A | Ribosome | 60S ribosomal protein L22-A |
| RPL8A | RL8A | Ribosome | 60S ribosomal protein L8-A |
| NOP58 | NOP58 | Ribosome | Nucleolar protein 58 |
| RPS1A | RS3A1 | Ribosome | 40S ribosomal protein S1-A |
| RPL26A | RL26A | Ribosome | 60S ribosomal protein L26-A |
| MAM33 | MAM33 | Miscellaneous | Mitochondrial acidic protein MAM33 |
| RPS23A | RS23A | Ribosome | 40S ribosomal protein S23-A |
| SSB1 | SSB1 | Ribosome | Ribosome-associated molecular chaperone SSB1 |
| RPS18A | RS18A | Ribosome | 40S ribosomal protein S18-A |
| RPL21A | RL21A | Ribosome | 60S ribosomal protein L21-A |
| HMT1 | HMT1 | Miscellaneous | Protein arginine N-methyltransferase 1 |
| RPL6A | RL6A | Ribosome | 60S ribosomal protein L6-A |
| PAB1 | PABP | Granule, stress granules | Polyadenylate-binding protein, cytoplasmic and nuclear |
| RPL18A | RL18A | Ribosome | 60S ribosomal protein L18-A |
| RPL3 | RL3 | Ribosome | 60S ribosomal protein L3 |
| NSR1 | NSR1 | Miscellaneous | Nuclear localization sequence-binding protein |
| RPS9A | RS9A | Ribosome | 40S ribosomal protein S9-A |
| STM1 | STM1 | Ribosome | Suppressor protein STM1 |
| ASC1 | GBLP | Ribosome | Guanine nucleotide-binding protein subunit beta-like protein |
| NOP1 | FBRL | Ribosome | rRNA 2'-O-methyltransferase fibrillarin |
| KRR1 | KRR1 | Ribosome | KRR1 small subunit processome component |
| TDH2 | G3P2 | Miscellaneous | Glyceraldehyde-3-phosphate dehydrogenase 2 |
| KRE33 | NAT10 | Ribosome | RNA cytidine acetyltransferase |
| CIC1 | CIC1 | Ribosome | Proteasome-interacting protein CIC1 |
| ENO2 | ENO2 | Miscellaneous | Enolase 2 |
| GBP2 | GBP2 | Miscellaneous | Single-strand telomeric DNA-binding protein GBP2 |
| FAS1 | FAS1 | Miscellaneous | Fatty acid synthase subunit beta |

| | | | |
|---|---|---|---|
| SHM1 | GLYM | Miscellaneous | Serine hydroxymethyltransferase, mitochondrial |
| TY1B-LR1 | YL11B | Miscellaneous | Transposon Ty1-LR1 Gag-Pol polyprotein |
| BFR2 | BFR2 | Ribosome | Protein BFR2 |
| CDC14 | CDC14 | Miscellaneous | Tyrosine-protein phosphatase CDC14 |
| YEF3 | EF3A | Granule, Ribonucleoprotein granules | Elongation factor 3A |
